# Dynamics of Bacterial and Viral Transmission in Experimental Microbiota Transplantation

**DOI:** 10.1101/2025.01.15.633206

**Authors:** James S. Weagley, Luis Alberto Chica Cárdenas, Ana Romani, Meagan E. Sullender, Somya Aggarwal, Heyde Makimaa, Michael P. Hogarty, Rachel Rodgers, Elizabeth A. Kennedy, Lynne Foster, Lawrence A. Schriefer, Megan T. Baldridge

## Abstract

Mouse models are vital tools for discerning the relative contributions of host and microbial genetics to disease, often requiring the transplantation of microbiota between different mouse strains. Transfer methods include antibiotic treatment of recipients and colonization using either co-housing with donors or the transplantation of fecal or cecal donor material. However, the efficiency and dynamics of these methods in reconstituting recipients with donor microbes is not well understood. We thus directly compared co-housing, fecal transplantation, and cecal transplantation methods. Donor mice from Taconic Biosciences, possessing distinct microbial communities, served as the microbial source for recipient mice from Jackson Laboratories, which were treated with antibiotics to disrupt their native microbiota. We monitored bacterial and viral populations longitudinally over the course of antibiotics treatment and reconstitution using 16S rRNA gene sequencing, quantitative PCR, and shotgun sequencing of viral-like particles. As expected, antibiotic treatment rapidly depleted microbial biomass and diversity, with slow and incomplete natural recovery of the microbiota in non-transplanted control mice. While all transfer methods reconstituted recipient mice with donor microbiota, co-housing achieved this more rapidly for both bacterial and viral communities. This study provides valuable insights into microbial transfer methods, enhancing reproducibility and informing best practices for microbiota transplantation in mouse models.

## Introduction

As appreciation grows for the diverse range of phenotypes the microbiota can mediate so does our reliance on manipulatable small animal models to experimentally interrogate these effects. The gut microbiota, a complex community of microorganisms residing in the gastrointestinal tract, plays a pivotal role in host health and physiology^1,2^. These microbial communities are involved in critical functions such as nutrient metabolism^3,4^, immune system development^5–7^, and protection against pathogens^8–10^. Dysbiosis, or the disruption of the normal microbiota balance, has been implicated in a range of diseases including obesity^11,12^, malnutrition^13,14^, and inflammatory bowel diseases^15,16^. Understanding the intricate relationships between the microbiota and host systems is therefore essential for developing therapeutic strategies targeting microbiota-related diseases.

To unravel these complex host-microbe interactions, researchers heavily leverage animal models, particularly mice. Controlled studies using mouse models enable the manipulation of microbial communities to observe resultant effects on host physiology and disease outcomes. A key aspect of such research is the generation of gnotobiotic mice— animals that are germ-free or colonized with specific, defined microbial communities^17^. These models provide a unique platform to study the microbiota’s role under controlled conditions. Despite the utility of germ-free mice for numerous applications, however, their use is associated with some drawbacks. For investigators studying numerous mouse genotypes, executing experiments not amenable to the use of a gnotobiotic isolator such as those involving particular pathogens or behavioral testing, or performing studies requiring a normal microbiota during development which is then disrupted, germ-free mice may be impractical or inappropriate. Antibiotics treatment represents an cost-effective and accessible alternative for the depletion of the microbiota, and this approach has been broadly used by many investigators to examine many of the same phenotypes explored in germ-free mice, often with strong concordance between the two models^17^.

Establishing gnotobiotic mice or reconstituting the microbiota after perturbation necessitates reliable and efficient methods for microbial transfer. Common techniques include fecal microbiota transplantation, cecal content transplantation, and co-housing. Fecal transplantation involves transferring fecal material from a donor to a recipient via the oral route, while cecal transplantation is similar but utilizes the cecal contents, which may have a different microbiota. Co-housing leverages the natural coprophagic behavior of mice to transfer microbes. Despite their widespread use, experiments designed utilizing these methods are generally based on historical precedent rather than a comprehensive understanding of their efficacy or the specific dynamics of microbial colonization they promote.

This reliance on convention can lead to assumptions that impact experimental outcomes and reproducibility, a major challenge facing the field^18–21^. For instance, variation in microbial transfer efficiency can affect the establishment of the desired microbiota, influencing the host’s physiological responses and potentially skewing results^22,23^. Furthermore, while bacterial community dynamics have been relatively well-studied, the transfer and establishment of viral communities (the virome) remain less understood. Given the growing recognition that bacteriophages and other viruses shape microbial communities and influence host health^24–30^, it is crucial to consider both bacterial and viral components when evaluating these methods. A comprehensive virome analysis is essential.

Herein, we systematically compare fecal transplantation, cecal transplantation, and co-housing as methods for transferring microbes to microbially depleted mice, characterizing this process longitudinally. Recipient mice from Jackson Laboratories were treated with antibiotics for one week to disrupt their native microbiota. Donor mice from Taconic Biosciences, known to have distinct microbial communities from Jackson, served as the source of bacteria and viruses for transplantation^31^. By employing 16S rRNA gene sequencing and quantitative PCR (qPCR), we monitored bacterial load, diversity, and composition. Additionally, we analyzed viral community composition and diversity through short-read shotgun sequencing of viral-like particles. We employed a comprehensive contig-based, annotation-agnostic strategy in conjunction with in-depth classification and binning of viral-like particle (VLP) sequences, allowing for an unbiased characterization of both known and novel viruses. Thus, we provide a more complete landscape of virome dynamics than would be possible with annotation-dependent methods alone.

We find that co-housing facilitates a more rapid transfer of both bacterial and viral communities compared to fecal and cecal transplants, which performed similarly to each other. Viral communities generally tracked with bacterial colonization, though each method enriched for distinct bacterial taxa—and their associated phages—at different timepoints. These results highlight method-specific influences on microbial community dynamics and underscore the importance of method selection in experimental design.

By carefully characterizing these commonly used methods, our study provides valuable insights that can enhance reproducibility and consistency in microbiota research. This work informs best practices for replacing the enteric microbiota of mice and contributes to our understanding of the mechanisms underlying microbial transfer and establishment in experimental models. Such insights are essential for advancing microbiota research and for the development of interventions targeting microbiota-related diseases.

## Results

### Effect of antibiotic treatment on bacterial load, diversity, and community composition

To evaluate the impact of antibiotic treatment on bacterial load, diversity, and community composition, we treated C57BL/6 mice from Jackson Laboratory (JAX) with a simplified antibiotic cocktail of vancomycin, neomycin, and ampicillin for seven days. This regimen, shown by us and others to mediate robust depletion of the microbiota within three to seven days while being well-tolerated by recipients^10,11^, was administered to JAX recipient mice. C57BL/6 mice from Taconic (TAC), which are well-established to harbor distinct enteric microbial communities compared to JAX mice^12,13^, served as microbiota donors, while untreated JAX and TAC mice served as controls (**Extended Data Fig. 1A**). Fecal samples were collected on days 0, 1, 3, and 7 for 16S rRNA gene qPCR and sequencing. At baseline (day 0), bacterial load was not significantly different between JAX and TAC mice ( **Extended Data Fig. 1B**, p > 0.05). By day 1, antibiotics significantly reduced bacterial load in JAX mice (p < 0.05), with 16S levels falling below the limit of detection by day 7 (**Extended Data Fig. 1B**). Bacterial load remained stable in untreated controls.

Sequencing of the V4 region of the 16S rRNA gene was performed and analyzed using the DADA2 pipeline implemented in Nephele to generate amplicon sequence variants (ASVs), which were grouped into bacterial species based on >97% sequence identity^32,33^. Initial ASV richness was comparable between JAX and TAC mice (mean ± SE: TAC = ∼80 ± 6 OTUs, JAX = 66 ± 5; **Extended Data Fig. 1C**). Antibiotic treatment significantly reduced species richness within one day compared to Jackson controls (mean ± SE: 43 ± 6 and 117 ± 2.4, respectively), with richness dropping to approximately one-sixth of the original level by day 7 (**Extended Data Fig. 1C**, 9.3 ± 0.9), although sequencing suggested some native bacteria persisted at low levels. To assess changes in community composition, unweighted UniFrac distances were calculated and visualized using principal coordinate analysis (PCoA). PCoA axis 1 explained 24.6% of the total variance, primarily reflecting antibiotic-induced shifts in JAX mice, while axis 2 (13.0%) separated untreated JAX and TAC samples (**Extended Data Fig. 1D**). Antibiotic treatment had a greater impact on microbial community composition than mouse vendor. These results demonstrate that one week of vancomycin, neomycin, and ampicillin treatment is sufficient to drastically reduce bacterial load and diversity and significantly alter gut microbiota composition.

### Route of transfer regulates community reconfiguration during transplantation following antibiotic treatment

We compared the outcomes of different transfer methods for transplanting TAC microbiota into antibiotic-treated JAX mice. Antibiotics were stopped on day 7, and mice were either maintained without intentional microbial transfer, gavaged on day 8 and 9 with prepared cecal material or fecal material or co-housed with untreated TAC donors for two weeks (**Fig. 1A**). TAC and JAX mice that didn’t receive antibiotics were also monitored over the same period.

**Figure 1.**
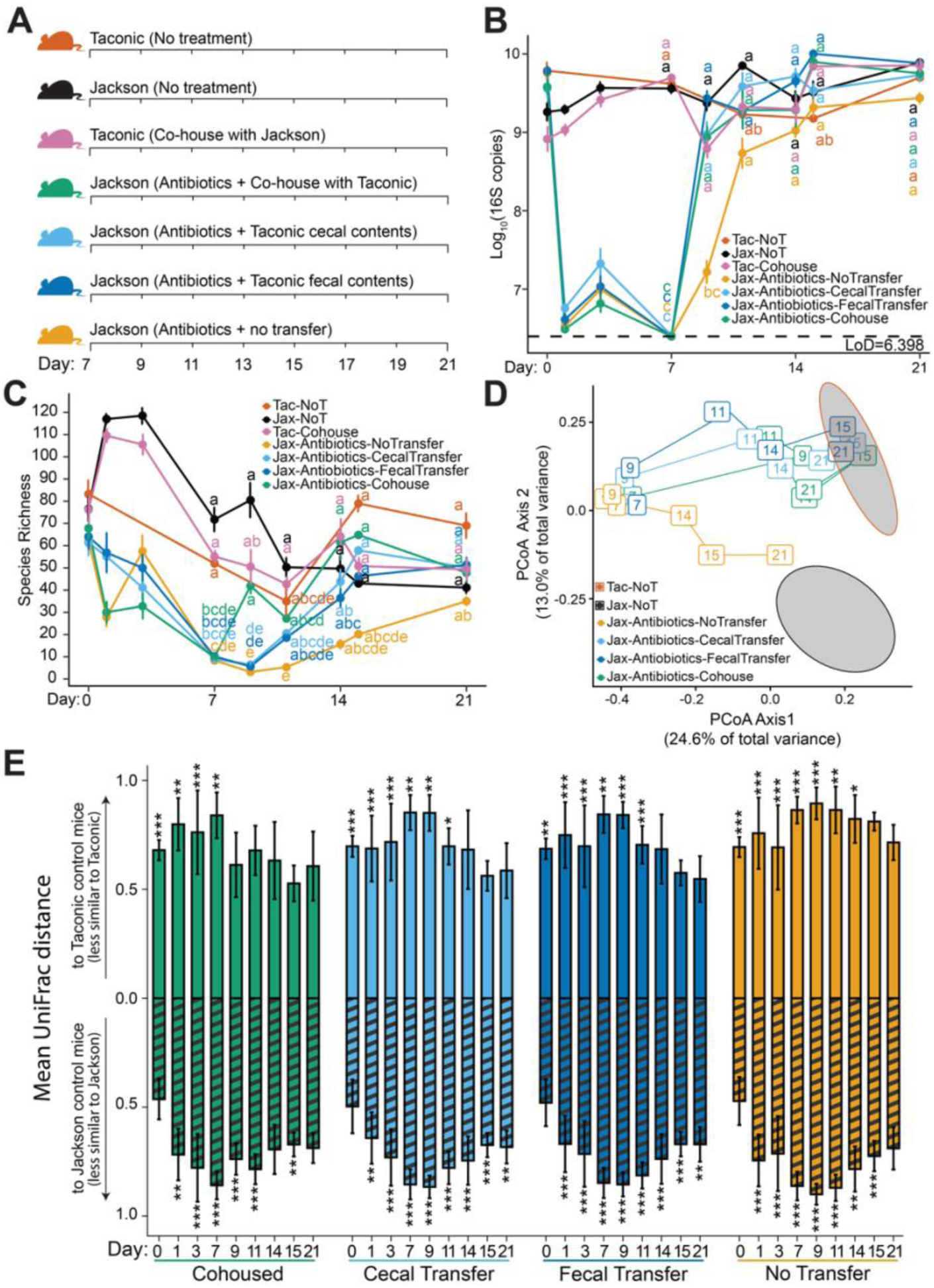
Cohousing of mice reconstitutes the bacterial microbiota more rapidly than fecal or cecal transplant. (**A**) Antibiotic-treated Jackson mice were either allowed to recover naturally (Jax-Antibiotics-NoTransfer; n=12, 8-12 samples per timepoint across three experiments), or were colonized with Taconic-associated microbes via gavage of cecal (Jax-Antibiotics-CecalTransfer; n=12, 7-12 samples per timepoint) or fecal (Jax-Antibiotics-FecalTransfer; n=12, 8-12 samples per timepoint) material, or co-housing (Jax-Antibiotics-Cohouse; n=11, 6-11 samples per timepoint). Jackson (Jax-NoT) and Taconic (Tac-NoT) mice that did not receive antibiotics were maintained as controls. (**B**) Bacterial load measured by qPCR of the 16S rRNA gene throughout the final two weeks of the experiment shows the rapid recovery of bacterial load following transplantation. The letters next to each point indicate significant groupings, those with the same letter were not significantly different from one another by two-way ANOVA followed by Tukey’s HSD post-hoc test for pairwise comparisons with an alpha of 0.05 (**C**) Bacterial species richness, the number of unique bacterial species observed per mouse, throughout the first week of the experiment via v4-16s sequencing. The letters next to each point indicate significant groupings, those with the same letter were not significantly different from one by Kruskal-Wallis followed by Bonferroni’s correction for multiple comparisons with an alpha threshold of 0.05. (**D**) Principal Coordinate Analysis of unweighted UniFrac distances calculated from species counts across all three weeks of the experiment illustrate the change in community composition following antibiotic treatment and microbiota transplantation. Ellipses (95% confidence) represent the baseline fecal bacterial communities of untreated Jackson (Jax-NoT) and Taconic (Tac-NoT) mice during the first week of the experiment. (**E**) Bar heights indicate the mean Unweighted UniFrac distance was calculated between the fecal bacterial community of Jackson mice treated with antibiotics that received microbial transplantation via co-housing, cecal transfer, fecal transfer, or no transfer, and the fecal bacterial community of control Jackson and Taconic mice that never received antibiotics, pooled by week (days pooled: [0,7), [7,14), [14,21]). Striped bars below the x-axis indicate the distance from untreated Jackson controls while solid bars above the x-axis indicate the distance from untreated Taconic controls. Error bars indicate the standard deviation of all pairwise comparisons between the indicated groups. A PERMANOVA test was performed on the unweighted UniFrac distance matrix for the comparisons shown followed by Bonferroni’s corrections for multiple comparisons (* *p* < 0.05, *\*\* p* < 0.01, \**\*\* p* < 0.001).

Bacterial load increased significantly by day 9 in all transplanted mice, regardless of method, compared to non-transplanted mice (**Fig. 1B**). While non-transplanted mice also showed detectable bacterial levels by day 9, their loads were nearly two orders of magnitude lower than those of transplanted or untreated control mice. By day 11, 16S copy numbers were comparable across all groups, and by day 14, bacterial loads remained similar for the rest of the experiment. In contrast, recovery of bacterial alpha-diversity progressed more slowly than bacterial load, particularly for the gavage based transplantation methods (**Fig. 1C**)

The effect of the transplant method on community composition was evaluated. Samples were separated by collection day and treatment group. Unweighted UniFrac distances assessed global changes in community composition and beta diversity. PCoA visualized microbial community recovery and transfer between days 7 and 21 (**Fig. 1D**). Confidence ellipses represented the original recipient (JAX) and donor (TAC) communities in feces from untreated mice from days 0 through 7.

Bacterial communities of antibiotic-treated JAX mice administered fecal or cecal transplants gradually approached the TAC donor community composition between days 9 and 11, converging by day 15 (**Fig. 1D**). While fecal or cecal transplant restored bacterial loads to TAC donor levels within two days (**Fig. 1B**), the community composition wasn’t fully recapitulated by day 9. In contrast, co-housed mice exhibited bacterial compositions equivalent to TAC donors after only two days (**Fig. 1D, E**). JAX mice that received antibiotics, but no transfer gradually returned towards untreated JAX mice, achieving a similar state to the original microbiota by day 21.

The unweighted UniFrac distances of treatment arms compared to untreated Taconic and Jackson control mice were analyzed using PERMANOVA. To account for temporal community composition drift and low sample sizes in untreated Taconic controls, daily treatment arm samples were compared to pooled control samples within the same week. Antibiotic-treated animals differed significantly from JAX controls after 1 day, with this difference persisting until day 7 (**Fig. 1E**). The bacterial community of all groups differed from TAC controls, increasing until day 7 (**Figs. 1D, E**). By day 9, antibiotic-treated JAX mice co-housed with TAC donors were no longer significantly different from TAC controls. Cecal or fecal transfer recipients differed from Taconic donors until day 14, suggesting varying transfer efficiency. Consistent with their bacterial loads (**Fig. 1B**), antibiotic-treated JAX mice recovered and were not significantly different from JAX controls by day 21 (**Figs. 1D, E**). However, the average distance between these groups remained elevated (mean ± SD: 0.69 ± 0.1) and hadn’t returned to baseline (mean ± SD: 0.47 ± 0.1). Co-housed mice showed the fastest transition to a TAC-like community but weren’t significantly different from JAX controls on days 14 and 21, though different on day 15. This may indicate incomplete microbial transfer or JAX bacteria recovery in co-housed mice, resulting in an ‘intermediate’ community state.

### Co-housing rapidly transfers Taconic-associated bacterial taxa and their predicted functional capabilities

To understand bacterial transfer following transplantation, ASVs were classified as TAC-(162 ASVs), JAX-associated (124 ASVs), or neither/both based on their prevalence in untreated mice in the samples collected during the first week of the experiment (50 TAC and 81 JAX samples) (**Fig. 2A**). These subsets were tracked throughout the experiment. TAC ASVs were higher in TAC mice than in untreated JAX mice, significantly so on days 7 and 8. Although there was a transient increase in the richness of TAC ASVs on days 1 and 3 in all groups measured, by day 7 the TAC-associated ASV richness in all JAX mice (mean = 0.7 – 2.6 Taconic ASVs per sample) was significantly lower than in TAC donors (∼36.3 Taconic ASVs per sample) (**Fig. 2B**). By day 9, TAC ASVs increased in co-housed mice (mean ± SE = 26.9 ± 2.0) and were higher than untreated JAX controls (mean ± SE = 0.7 ± 0.1) but not significantly different from TAC co-housed donors (mean ± SE = 30.8 ± 4.5). At day 9, cecal or fecal transplants didn’t significantly increase TAC-associated ASV richness (means ± SEs = 4.6 ± 0.8 and 4.3 ± 0.4, respectively). By day 15, the richness of TAC-associated ASVs in JAX mice that received antibiotics followed by cecal or fecal transplant was significantly elevated relative to Jackson control mice and remained relatively steady until the end of the experiment on day 21. Co-housing introduced donor-specific bacteria more rapidly than cecal or fecal transplant.

**Figure 2.**
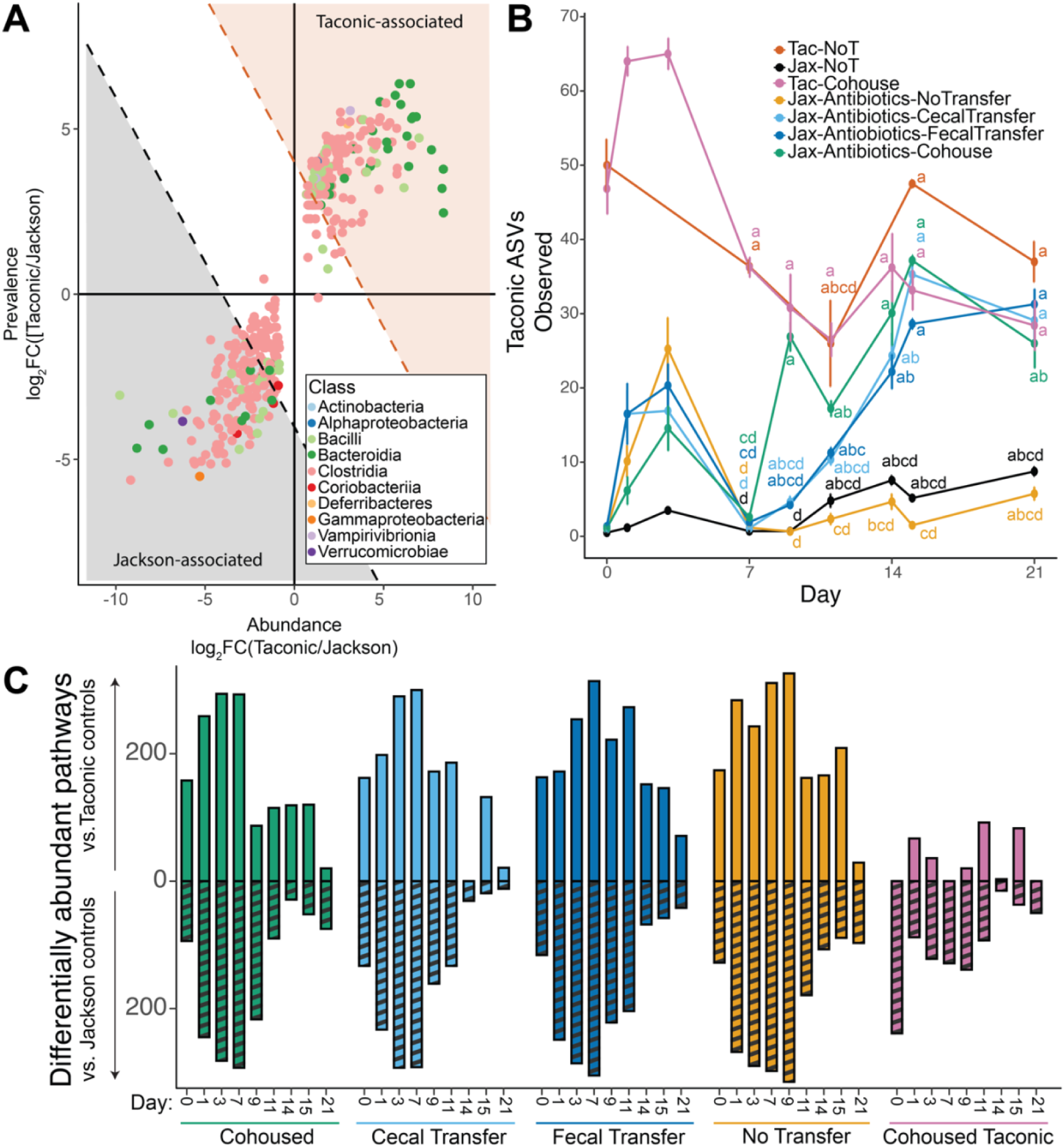
Defining and tracking Taconic-associated ASVs and their predicted functional capacities. (**A**) Bacterial ASVs were defined as being Taconic or Jackson associated based on their prevalence and abundance in samples collected from antibiotics-free JAX (n = 81) and TAC (n=50) mice over the first week of the experiment. ASVs that had an absolute combined log_2_(Fold Change) greater than 4 were consider Taconic-associated while those that had an absolute combined log_2_(Fold Change) less than −4 were considered Jackson-associated. (**B**) Richness of Taconic ASVs was monitored throughout the course of the experiment, plot show mean richness in each group at a given timepoint, error bars represent the standard error, compact letter display indicates results of pairwise Kruskal-Wallis followed by Bonferroni’s correction for multiple comparisons with an alpha threshold of 0.05. (**C**) Abundance of functional pathways were predicted using PICRUSt2. Bar heights indicate the number of pathways determined to be significantly differentially abundant using DESeq2 (alpha = 0.05) to compare between the fecal bacterial community of Jackson mice treated with antibiotics that received microbial transplantation via co-housing, cecal transfer, fecal transfer, or no transfer, and Taconic co-housed mice, and the fecal bacterial community of control Jackson and Taconic mice that never received antibiotics, pooled by week (days pooled: [0,7), [7,14), [14,21]).

To characterize the functional capacity of bacterial communities, we used PICRUSt2 to predict the abundance of MetaCyc pathways in each sample^34,35^. We compared antibiotic-treated JAX mice and TAC co-housed donors to JAX and TAC controls grouped by week to identify differentially abundant pathways using DESeq2 (**Fig. 2C**)^36^. A larger number of differentially abundant pathways indicates greater functional disparity. Consistent with our taxonomic results, the number of differentially abundant pathways between co-housed JAX mice and TAC control mice dropped off rapidly, from over 200 on day 7 to less than 100 on day 9. Mice receiving a cecal transplant didn’t drop below 100 until day 14, while fecal transplant recipients didn’t until day 21. Co-housing led to a more rapid and comprehensive transfer of Taconic-associated bacterial taxa and their predicted functional capabilities compared to cecal or fecal transplantation.

### Transfer of specific bacterial taxa differs between methods of transplantation

Changes in bacterial community composition were profiled throughout the experiment. The microbiota of control communities remained stable (**Fig. 3A**). The relative abundance of *Bacilli* and *Gammaproteobacteria* increased in mice receiving the antibiotic cocktail by day 3 (**Fig. 3A**), though the overall bacterial load and richness remained low (**Fig. 1B,C**). By day 9, co-housed mice were significantly enriched for TAC ASVs from the *Tannerellaceae*, *Prevotellaceae*, and *Muribaculaceae* families, and *Lactobacillaceae* ASVs that were not associated with either supplier, relative to mice without transplantation (**Fig. 3B**). Mice receiving cecal or fecal transplants were also enriched for these *Lactobacillaceae* ASVs, along with un-associated *Enterobacteriaceae* ASVs (**Fig. 3B**). This may reflect increased aerotolerance and colonization of uninhabited gastrointestinal tract niches. *Bacteroidaceae* ASVs associated with the original TAC communities were enriched in mice receiving fecal or cecal transplants (**Fig. 3B**). *Enterobacteriaceae* ASVs were significantly enriched in mice receiving cecal transplant compared to co-housed mice, while co-housed mice were enriched for *Muribaculaceae* ASVs.

**Figure 3.**
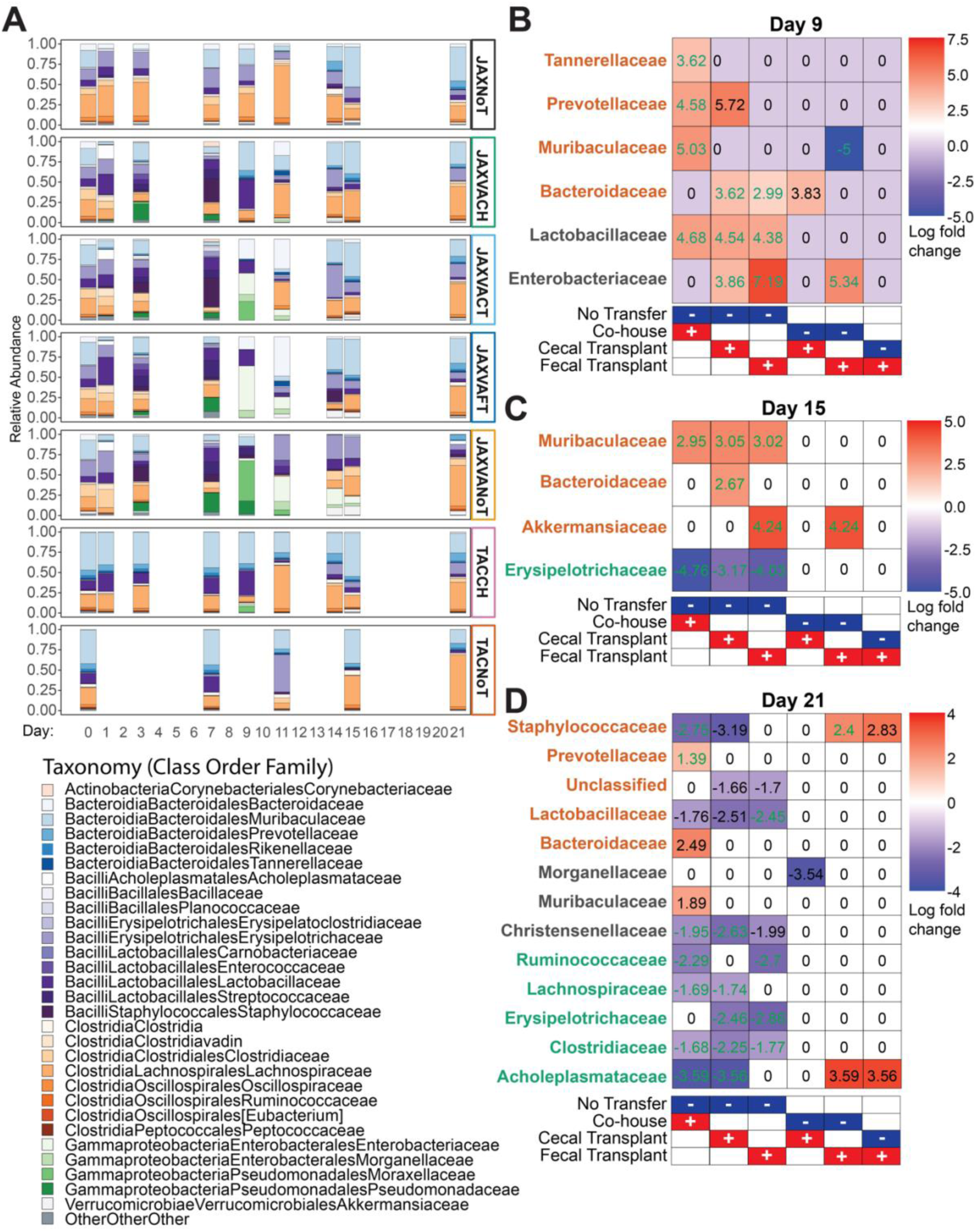
Specific bacterial taxa are enriched across transplantation methods. **(A**) Mean relative abundance of bacterial families throughout the course of the experiment in each of the treatment groups, averaged across all samples on each day. (**B-D**) Bacterial families associated with original bacterial communities of mice from Taconic (orange), Jackson (green) or neither/both (black) that were significantly enriched or depleted for the comparisons shown below each plot as determined using ANCOM-BC2. Panels show results from days 9 **(B)**, 15 **(C)**, and 21 **(D)**. Log fold change values in green remained significant after applying a stringent mixed directional FDR control.

TAC-associated ASVs from the *Muribaculaceae* family were significantly enriched in all mice receiving a microbiota transfer at day 15, regardless of method (**Fig. 3C**). TAC *Bacteroidaceae* ASVs were enriched in mice receiving a cecal transplant, while TAC-associated *Akkermansiaceae* ASVs were enriched in mice receiving a fecal transplant compared to the ‘no transfer’ and co-housed groups. JAX-associated *Erysipelotrichaceae* ASVs were the first to recover in antibiotic-treated mice without a transplant and were significantly enriched by day 15 compared to co-housed mice or mice receiving a fecal or cecal transplant (**Fig. 3C**).

By day 21, significant differences in bacterial taxa enrichment were observed across groups (**Fig. 3D**). TAC *Staphylococcaceae* ASVs were enriched in the ‘no transfer’ and fecal transplant groups compared to co-housed and cecal transplant recipient mice, while TAC *Lactobacillaceae* ASVs were enriched in the ‘no transfer’ group compared to the transplant groups. Notably, *Christensenellaceae* ASVs, not associated with either supplier, were enriched in the ‘no transfer’ group compared to all transplantation groups. Several JAX-associated taxa, including *Ruminococcaceae*, *Lachnospiraceae*, *Erysipelotrichaceae*, and *Clostridiaceae*, were enriched in the ‘no transfer’ group compared to all transplant groups. The fecal transplant groups exhibited distinct enrichment patterns, including higher levels of TAC-associated *Staphylococcaceae* and JAX-associated *Acholeplasmataceae* compared to co-housed or cecal transplant recipient mice. JAX *Acholeplasmataceae* were also enriched in the ‘no transfer’ group compared to co-housed or cecal transplant recipient mice. Distinct transplantation methods differentially enriched specific bacterial taxa over time, with co-housing favoring *Muribaculaceae* and certain donor-associated lineages earlier, while fecal and cecal transplants more strongly enriched for *Bacteroidaceae*, *Enterobacteriaceae*, and *Akkermansiaceae*; mice without transplants exhibited recovery of native JAX-associated taxa and unique enrichments not observed in other groups by day 21.

### Bacterial diversity and community composition differ across gastrointestinal sites but not the method of transfer

Bacterial diversity and community composition varied across gastrointestinal sites and treatment groups. Lower richness was consistently observed in ileal contents compared to cecal contents and stool, as well as at all sites in mice without microbiota transfer after antibiotic treatment (**Fig. 4A**). Stool richness trended higher in transplanted mice than JAX controls but lower than TAC controls, suggesting incomplete bacterial taxa transfer. PCoA illustrated community composition differences, with samples clustering according to treatment and site (**Fig. 4B**). Samples from JAX mice treated with antibiotics but no transfer clustered together, away from samples from transplanted mice and control communities, with samples from the same gastrointestinal site clustering together. To compare community compositions across treatments, PCoA was performed on samples subset by the gastrointestinal site (**Figs. 4C-E**). Ileal, cecal, and stool communities of JAX co-housed mice were closely aligned with TAC mice, while ileal communities of JAX fecal or cecal transplantation mice clustered separately (**Fig. 4C**), a difference that was not as pronounced in cecal or stool samples (**Figs. 4D, E**). These findings provide insights into the dynamics of bacterial community re-establishment across microbiota transplantation methods.

**Figure 4.**
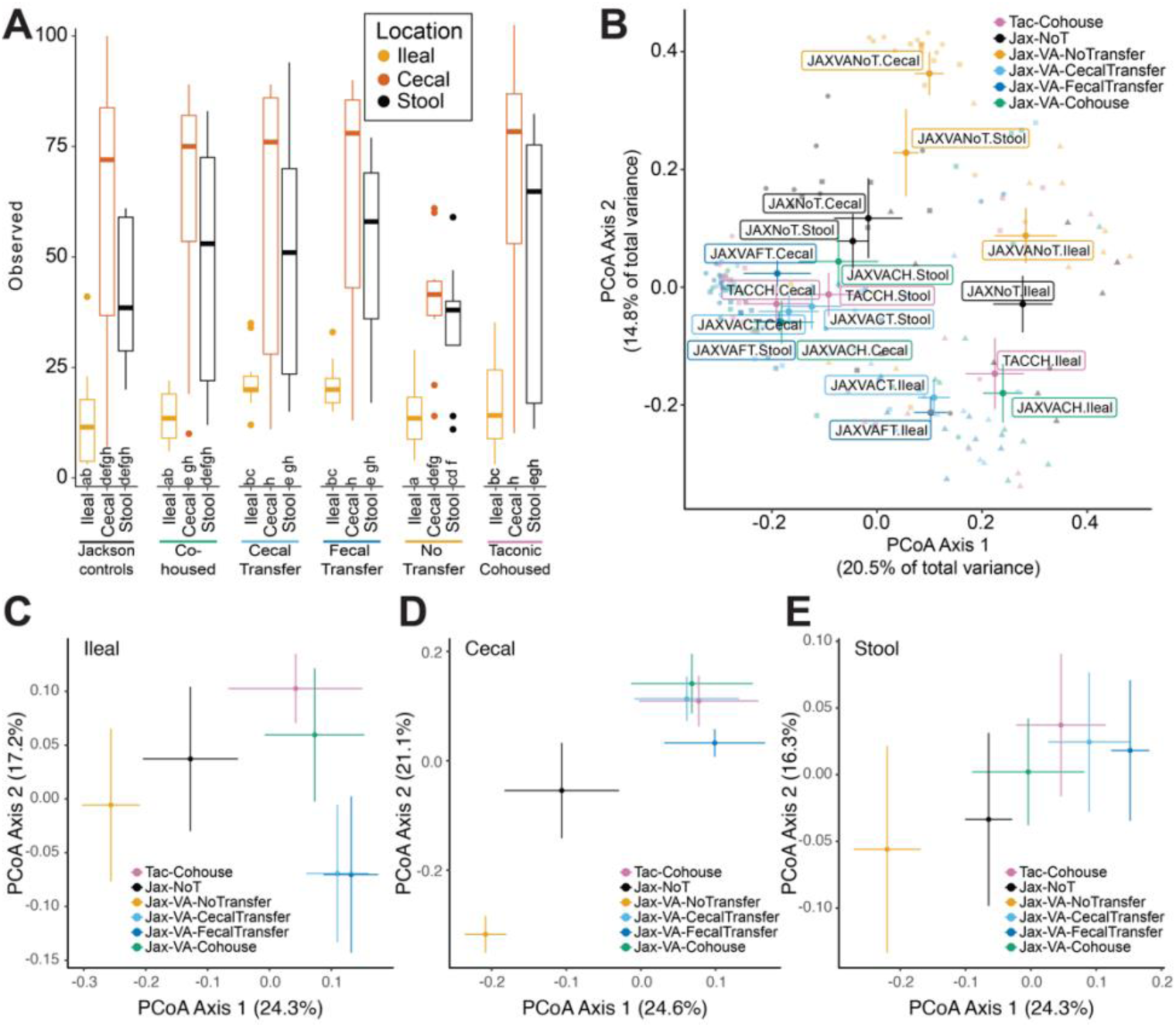
Bacterial diversity and community composition across different body sites and treatment groups. (**A**) Bacterial richness, the number of bacterial taxa observed in cecal, ileal, and stool samples across different groups, including Jackson controls, co-housed, cecal transfer, fecal transfer, no transfer, and Taconic controls. The letters under each bar indicate significant groupings, those with the same letter were not significantly different from one another. Observed richness was compared using a linear mixed-effects model with sequencing library as a random effect, post hoc comparison of marginal means were Tukey-adjusted; significance groups are indicated by differing letters (α = 0.05). (**B**) Plot of Principal Coordinate Analysis (PCoA) of Unweighted UniFrac Distances illustrating bacterial community composition from cecal, ileal, and stool samples, across each method of transplantation, including Jackson controls and cohoused Taconic donors. PCoA of Unweighted UniFrac Distances of samples on Day 21 across treatment groups from (**C**) ileal contents, (**D**) cecal contents, or (**E**) stool.

### Viral Community is Depleted Post-Antibiotics

We examined the viral response to antibiotic treatment, noting a significant reduction in bacterial load and diversity over the first week (**Figs. 1B, C**). Short-read shotgun sequencing of DNA from viral-like particles (VLPs) assessed the impact of antibiotics on viral communities. Viral richness (unique contigs from sequencing data) was compared from days 0 to 7 between TAC and JAX controls and antibiotic-treated JAX mice treated. Read counts were normalized to reduce library batch effects. Viral richness dropped significantly from 596 ± 84 (mean ± SE) unique viral contigs on day 0 to 34 ± 2 on day 7 (**Extended Data Fig. 2A**). This pattern persisted when contigs with clear viral signatures were analyzed and when aggregated into clusters representing viral species, genera, and families (**Extended Data Fig. 2A**). The reduction in viral richness indicates viral populations’ susceptibility to bacterial community disruptions, likely due to their dependence on bacterial hosts. A notable shift in viral community composition, primarily driven by antibiotic treatment, was observed in PCoA of Jaccard distances (**Extended Data Fig. 2B**). The separation along axis 1 was mainly between antibiotic-treated and untreated groups, highlighting antibiotics’ substantial impact on viral community composition. Similar results were found when considering only high-quality viral contigs or different taxonomic ranks (**Extended Data Fig. 2B**).

### Co-housing facilitates more rapid transfer of donor viral communities

To evaluate the effects of different microbiota transfer methods on viral communities in antibiotic-treated mice, we compared viral richness and community composition post-transplantation. Samples were collected from days 7 to 14 to observe viral population reestablishment. This analysis included all contigs from short sequencing of VLPs, avoiding biases from viral annotation and prediction algorithms. Viral richness, measured by unique viral contigs, did not uniformly increase across all groups after antibiotics ceased. By day 9, co-housed JAX mice showed a quicker rise in viral richness than other groups, with no significant difference from TAC mice (**Fig. 5A**). Viral richness was low in mice with fecal or cecal transfers, or no transfer, at this point. By day 11, viral richness increased in all groups, and while transplantation groups had lower richness than Taconic donors, the difference was not significant. Antibiotic-treated mice without transplants had significantly lower viral richness, which only slightly increased by day 11. By day 14, viral richness in all transfer groups converged, similar to untreated controls, indicating successful re-colonization.

**Figure 5.**
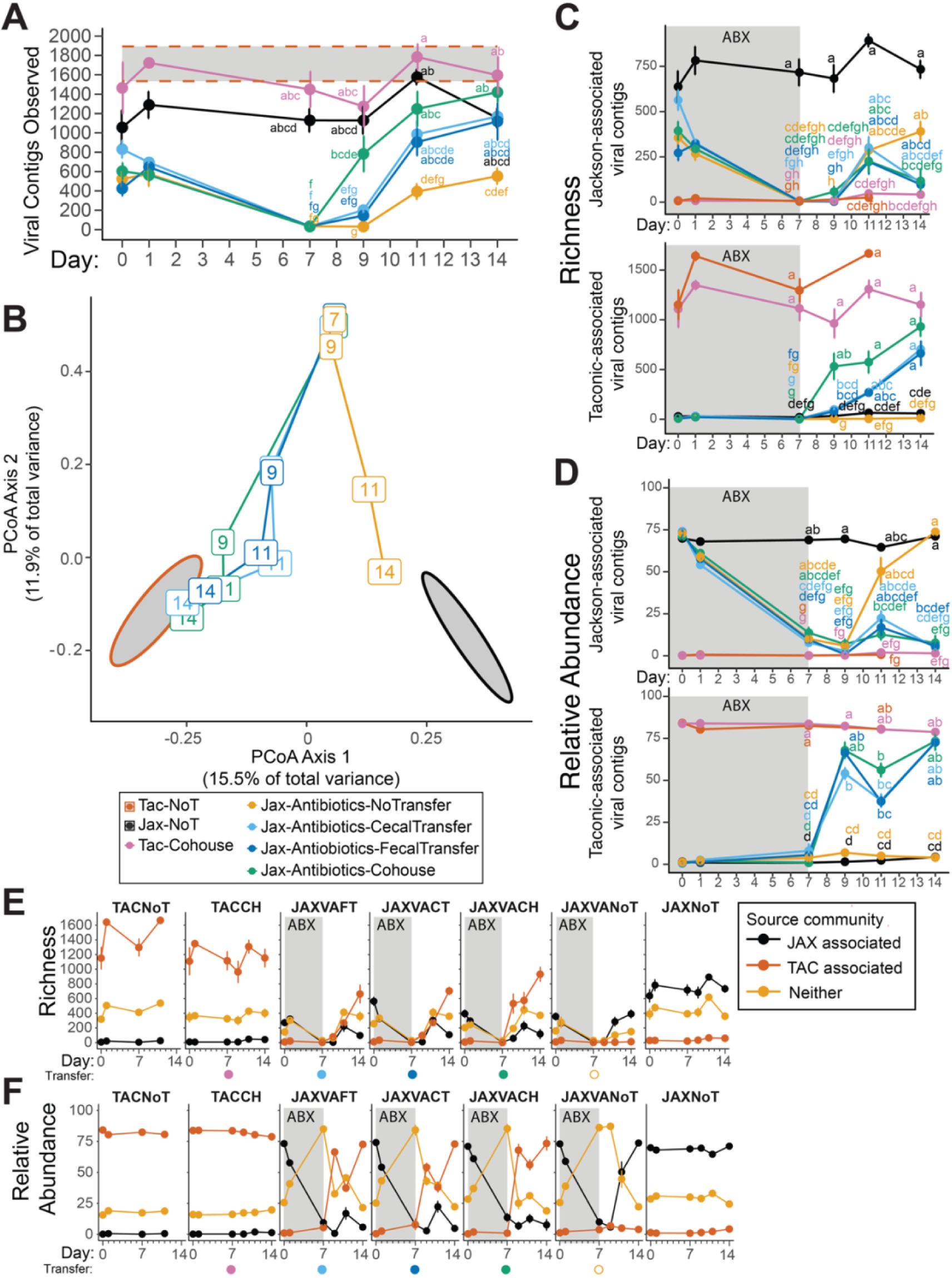
Cohousing of mice reconstitutes the viral microbiota more rapidly than fecal or cecal transplant. (**A)** Viral richness from days 7 through 14 of the experiment shows the rapid recovery of viral diversity following transplantation. The letters next to each point indicate significant groupings, those with the same letter were not significantly different from one another by pairwise Kruskal-Wallis followed by Bonferroni’s correction for multiple comparisons with an alpha threshold of 0.05 (**B**) Principal Coordinate Analysis of Jaccard distances calculated from contig read counts across two weeks of the experiment illustrate the change in community composition following antibiotic treatment and microbiota transplantation. Ellipses (95% confidence) represent the baseline fecal viral communities of untreated Jackson (Jax-NoT) and Taconic (Tac-NoT) mice. (**C**) Richness and (**D**) relative abundance of viral contigs defined as (i) Jackson or (ii) Taconic-associated throughout the course of the experiment. The letters next to each point indicate significant groupings, those with the same letter were not significantly different from one by pairwise Kruskal-Wallis followed by Bonferroni’s correction for multiple comparisons with an alpha threshold of 0.05. The (**E**) richness and (**F**) relative abundance of viral contigs associated with mice from Jackson, Taconic, or neither/both, over the first two weeks of the experiment. n = 6-11 samples per treatment group and timepoint, for all groups excluding TACNoT controls (n = 1-5)

We assessed the impact of transplantation methods on viral community composition using PCoA of Jaccard distances. Results indicated a gradual alignment of viral communities in fecal and cecal transplanted mice with TAC donor profiles between days 9 and 11, achieving full convergence by day 14 (**Fig. 5B**). Co-housed mice showed rapid convergence with donor profiles by day 9, suggesting co-housing enhances efficient viral transfer. In contrast, antibiotic-treated mice without transplants experienced delayed recovery, remaining distinct from donor and control profiles until the final sampling day, gradually nearing the original JAX community by day 14.

These findings highlight the critical role of microbiota transplantation in reestablishing viral communities. Co-housing was the most effective for rapid and complete viral transfer, while fecal and cecal transplants also supported successful colonization, though more slowly. Mice without transplants showed delayed and incomplete viral community recovery, emphasizing the need for external microbial inputs to restore gut virome diversity and composition post-antibiotic treatment.

### Transfer of Taconic-associated viruses to Jackson mice

To monitor virus movement between TAC and JAX mice, viral contigs were categorized as TAC, JAX, or neither/both, based on their prevalence and abundance in samples from the first week of the experiment (34 TAC and 50 JAX samples) without antibiotics. Out of 6,249 contigs, 2,535 were TAC-associated and 1,129 were JAX-associated. Antibiotic treatment significantly reduced the richness and relative abundance of JAX-associated viruses (**Figs. 5C-F**). TAC and JAX control mice maintained consistent viral richness and abundance (**Figs. 5C-F**), with TAC mice having more unique viral genome contigs than JAX controls (**Figs. 5C, E**).

After microbiota transplantation, TAC-associated virus richness and abundance in co-housed JAX mice was not significantly different from TAC donors by day 9 (**Figs. 5C, D**). By day 9, TAC-associated virus relative abundance in fecal and cecal transplant recipients was similar to donors (**Fig. 5D**), though diversity was lower (**Fig. 5C**). By day 11, four days after transplantation, TAC-associated virus abundance in fecal or cecal transplant recipients dropped well below donor levels (**Figs. 5D, F**). In contrast, co-housed mice maintained relatively stable TAC-associated virus abundance. This decline was accompanied by a slight resurgence of JAX-associated viruses on day 11 (**Figs. 5D, F**), but this rebound was not sustained, except in JAX antibiotic-treated mice without transplantation, where JAX-associated viruses persisted (**Figs. 5D, F**).

Despite only receiving two gavages of fecal or cecal contents, the number of unique viral genome fragments in transplanted mice increased, with detectable TAC-associated virus levels rising even after one week (**Figs. 5C, E**). However, TAC-associated virus richness in JAX mice with cecal or fecal transplants remained significantly different from TAC donors one week after gavage, on day 14 (**Figs. 5C, E**).

Viral diversity in JAX mice treated with antibiotics and no transplantation remained depleted throughout the experiment (through day 14), despite JAX-associated virus abundance returning to untreated JAX control and day 0 levels by day 11 (**Figs. 5C-F**). This indicates a long-lasting impact of antibiotics on viral diversity in mice without microbial transplants, despite partial recovery of JAX-associated viral abundance.

These findings underscore the complexity of viral community dynamics post-microbiota transplantation, with co-housing being the most effective method for rapidly restoring donor-associated viral communities. While fecal and cecal transplants also facilitated donor virus colonization, the process was slower and less consistent when considering viral richness and abundance.

### Binning viral genomes confirms rapid microbial transfer following cohousing

Contigs were binned into homogeneous groups representing individual viral genomes using CoCoNet^37^, based on sequence composition and coverage variation. 1,525 contigs were assigned to 629 unique bins (mean 2.4 contigs per bin, 426 bins with only one contig). Bins were categorized as JAX- or TAC-associated, or neither/both, according to previous contig classifications. No bins contained both JAX and TAC-associated contigs, allowing entire bins to be classified based on the presence of any TAC- or JAX-associated contigs (**Figs. 6A-E**). The richness of JAX-associated bins significantly decreased by day 7 in antibiotic-treated mice (**Fig. 6B**). The richness of TAC-associated bins increased significantly on day 9 in co-housed mice (**Figs. 6A, C**), but did not reach similar levels until day 14 in fecal or cecal transplant mice. Additionally, the average abundance of bins was quantified based on sequencing read depth, normalized by bin length and sample sequencing depth. The average abundance of JAX bins (RPKM) slightly decreased after antibiotic treatment and was significantly lower on day 9 in mice with microbiota transplantation (**Fig. 6D**). The average abundance of TAC bins sharply increased in all transplant mice on day 9 (**Fig. 6E**). These findings align with our annotation free contig-level analysis, showing a rapid increase in viral richness in co-housed mice compared to those with fecal or cecal transplants (**Figs. 5C, 6C**). Meanwhile, the average abundance (RPKM) of Taconic bins increased in all transplant groups by day 9, mirroring the increased relative abundance observed in the contig-based analysis (**Figs. 5D, 6E**).

**Figure 6.**
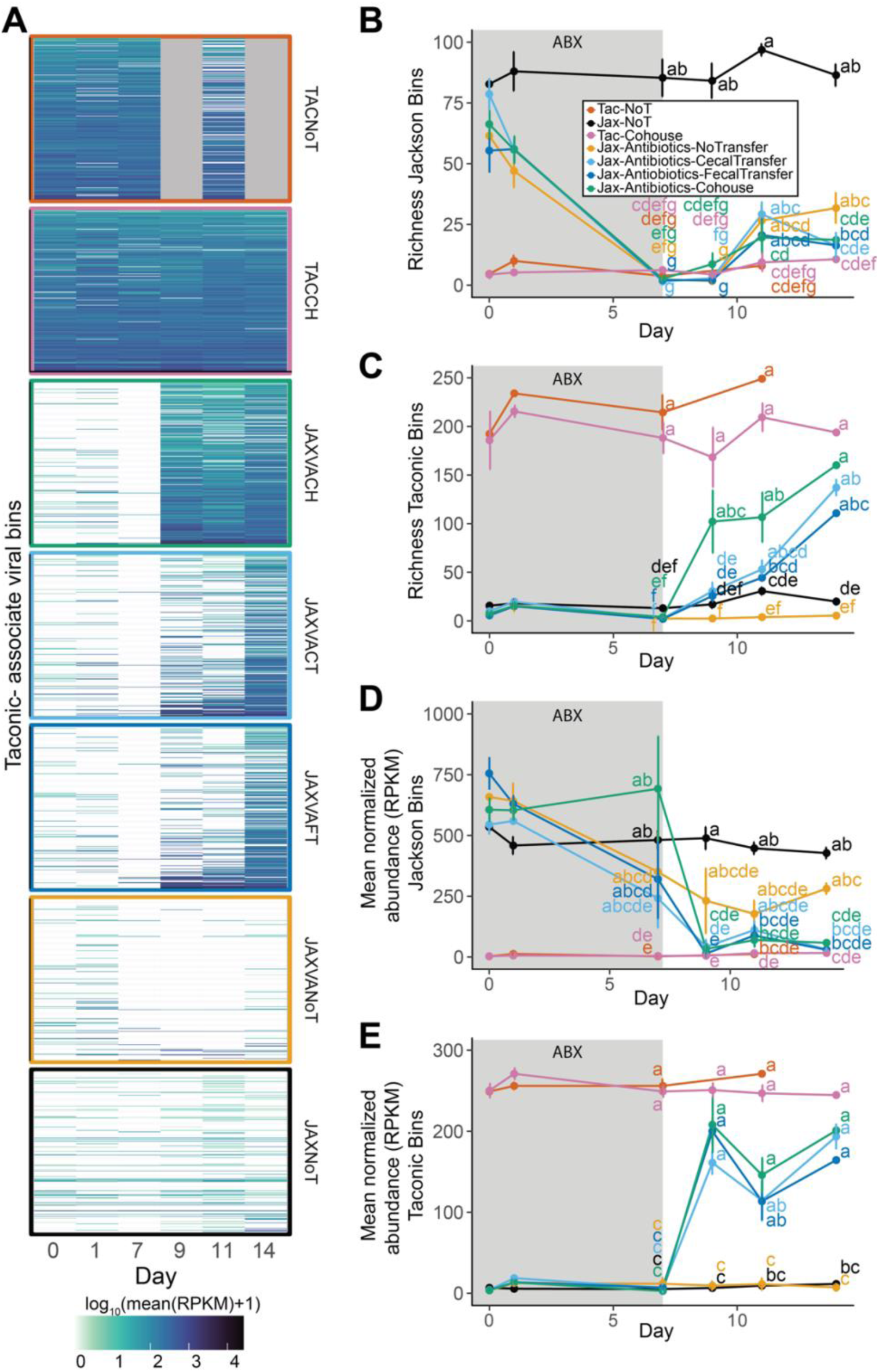
Binning viral genomes confirms rapid microbial transfer following cohousing. (**A)** Heatmap illustrating the richness of Taconic-associated viral bins over time in each treatment and control group as indicated on the right side of each panel. Line plots of the richness of (**B**) Jackson- and (**C**) Taconic-associated viral bins and mean normalized abundance of (**D**) Jackson- and (**E**) Taconic-associated viral bins over the course of the experiment. The letters next to each point indicate significant groupings, Those with the same letter were not significantly different from one another by Kruskal-Wallis test followed by Bonferroni correction for multiple comparisons with an alpha of 0.05. n = 6-11 samples per treatment group and timepoint, for all groups excluding TACNoT controls (n = 1-5)

### Dynamics of bacteriophages predicted to target discriminatory bacterial taxa

We examined the dynamics of phages targeting bacterial families with differential abundance during transplantation. *Tannerellaceae*, *Prevotellaceae*, *Muribaculaceae*, *Bacteroidaceae*, *Lactobacillaceae*, *Enterobacteriaceae*, *Akkermansiaceae*, and *Erysipelotrichaceae* distinguished treatment groups at days 9 and 15 (**Figs. 3B, C**). Using iPHoP^38^, we assessed the predicted bacterial host range for filtered phage contigs. No phages were predicted to target *Erysipelotrichaceae* or *Prevotellaceae*, and *Akkermansiaceae*-targeting phages were rare and not significantly different across groups. The abundance of phages targeting the remaining bacterial families varied over time (**Fig. 7A**). Phages targeting *Bacteroidaceae* were elevated on day 9 in cecal and fecal transplant recipients, aligning with bacterial data (**Fig. 3B**). Most phages in transplant recipients were temperate and not specifically linked to the original JAX or TAC communities (**Fig. 7B**), with a few originally TAC-associated. JAX controls maintained JAX-associated phages targeting *Bacteroidaceae*, predicted to be virulent. Also mirroring the abundance of their bacterial hosts, *Enterobacteriaceae*-targeting phages peaked on day 9 in fecal transplant recipients (**Figs. 7A, B**). All *Lactobacillaceae*-hosted phages were temperate and increased on day 9 in transplant groups (**Figs. 7A, B**). *Muribaculaceae*-hosted phages were not temperate, with elevated abundance on day 9 in co-housed mice (**Figs. 7A, B**). Although many *Muribaculaceae*-targeting phages were predicted to be virulent, their abundance still matched their bacterial hosts (**Fig. 3B**). Despite *Tannerellaceae* bacterial enrichment on day 9 in co-housed mice, their targeting phages were not elevated. Phage abundance closely mirrored their predicted bacterial hosts, suggesting induction of temperate phages post fecal or cecal transplant, in contrast to transfer of virulent phage specifically targeting a high-abundance bacterial community member, *Muribaculaceae,* in the context of co-housing.

**Figure 7.**
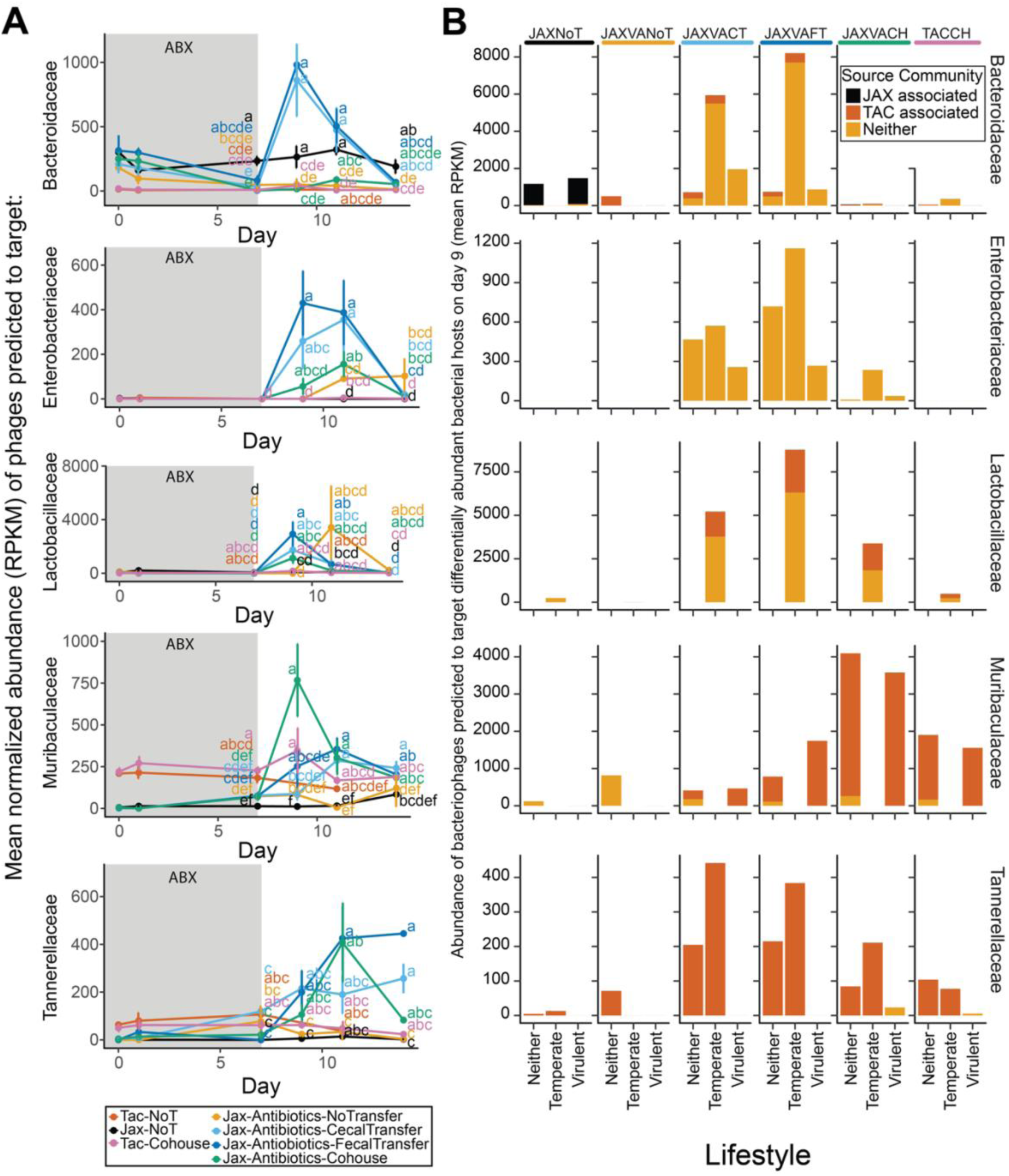
Abundance of bacteriophages mirrors discriminatory bacterial hosts. (**A**) The mean normalized Reads Per Kilobase per Million mapped reads (RPKM) abundance of phage bins predicted to target bacteria that were differentially abundant on day 9 (Fig. 3B) or day 15 (Fig. 3C), over time. Other bacterial families were either not predicted to be hosts for phages in this dataset, or were not significantly different between any of the groups at the timepoints test (Akkermansiaceae). The letters next to each point indicate significant groupings, those with the same letter were not significantly different from one another by Kruskal-Wallis test followed by Bonferroni correction for multiple comparisons with an alpha of 0.05. (**B**) Bar plots illustrating the abundance (mean RPKM) of phage bins targeting discriminatory bacterial taxa on day 9. Each column represents a treatment group. Within each plot, phages are split based on their predicted lifestyle. Rows are faceted by the family of the predicted bacterial host. n = 6-11 samples per treatment group and timepoint, for all groups excluding TACNoT controls (n = 1-5)

## Discussion

This study compared co-housing, fecal, and cecal transplantation in antibiotic-treated mice to reconstitute bacterial and viral communities^22,23,31,39^. Transplanting donor microbiota allows researchers to examine how composition influences health and disease. However, there’s a lack of systematic comparisons of commonly used transfer methods. We addressed this gap by assessing the efficiency, dynamics, and ecological outcomes of three widely used techniques. Our findings are relevant for improving experimental methods and enhancing the reproducibility of mouse-based studies on host-microbe interactions.

Our study indicates that co-housing is the most effective method for rapid microbiota transfer, quickly reconstituting both bacterial and viral communities. Sustained microbial exchange through natural behaviors like coprophagy and direct contact mirrors ecological interactions in natural settings. This efficiency aligns with previous studies showing that co-housing facilitates the transfer of donor-associated microbial taxa more robustly than gavage methods^31,40^. Co-housed mice rapidly reconstitute microbial diversity and functionality, making it useful for time-sensitive microbiota establishment. Fecal and cecal transplantation, though less efficient in the short term, achieved comparable outcomes by day 21. These methods may still be valuable in experimental scenarios where co-housing is impractical, but the slower pace of microbial establishment emphasizes the importance of considering time frames in experiments. Researchers should select transplantation methods that align with study goals to optimize reproducibility and minimize variability.

Antibiotic cocktail treatment significantly reduced intestinal bacterial loads within a day, suggesting that shorter treatment duration from the currently commonly-used 1-2 weeks before microbiota transfer would still be effective. Mice without transplantation showed delayed and incomplete microbiota recovery. This limited natural recolonization indicates insufficient native microbial recovery to restore pre-treatment diversity within two weeks, but does highlight potential for gut microbiota resiliency over the longer term.

The study highlighted the interdependence of bacterial and viral dynamics, reflecting the ecological relationships between bacteriophages and their hosts. Antibiotics rapidly depleted bacteria, leading to a slightly delayed but robust reduction in phage communities. Co-housed mice quickly established donor-associated bacterial taxa like *Muribaculaceae* and *Tannerellaceae*, along with their phages. In contrast, early introduction of bacteria such as *Bacteroidaceae*, *Lactobacillaceae*, and *Enterobacteriaceae* via fecal or cecal transplant, was linked to temperate phages targeting these hosts, which were not always main members of the donor’s viral community. These phages likely emerged via induction from donor pioneering bacteria during early transplantation. The synchronized reconstitution seen with co-housing emphasizes the role of ecological interactions in microbial recovery, suggesting that co - housing may offer a more natural environment for community establishment.

The site-specific variation in microbial recovery across gastrointestinal regions underscores the complexity of microbiota reconstitution. The distinct microbial diversity and composition between ileal, cecal, and fecal samples emphasize the importance of sampling multiple regions to capture microbiota dynamics. These findings align with previous work showing anatomical differences in microbial communities and their functions ^41–44^.

Systematic evaluations of transplantation methods across different contexts are crucial. Future studies should investigate how donor-recipient compatibility, microbial complexity, and host genotype influence outcomes. While this study focused on bacterial and viral communities, incorporating fungi and archaea will provide a more comprehensive understanding. Advancements in functional and longitudinal metagenomic analyses, coupled with improved animal models, will enhance our ability to study microbiota establishment and its implications for host health and disease. As we expand comparisons to new donor/recipient sources and microbial compartments, our comprehensive virome profiling pipeline, utilizing contig-based, classification-agnostic, and robust binning and classification methods, will ensure consistent capture of even poorly characterized or emerging viral populations across different experimental models. While characterization of human viral communities has broadly lagged behind the study of bacterial communities, the virome of experimental mice remains even more enigmatic. Our method successfully facilitated virome analysis and comparisons, and it could also be extended to analyze other poorly annotated microbes like fungi and archaea.

This study systematically compared microbiota transplantation methods, refining experimental approaches in microbiome research and revealing ecological and functional dynamics underlying microbial reconstitution. These insights will facilitate tailoring of transplantation methods to individual studies, thereby enhancing reproducibility and translational potential, and advancing our understanding of the microbiome’s role in health and disease.

## Materials and Methods

### Mice and Treatments

WT C57BL/6J (stock no. 000664) mice were purchased from Jackson Laboratories (JAX) and C57BL/6N (stock #B6), murine pathogen free health standard, mice were purchased from Taconic Biosciences (TAC) and maintained at Washington University School of Medicine under specific-pathogen-free conditions according to University guidelines. Female mice, aged six to eight weeks, were exclusively used in all experiments to facilitate cohousing experiments. Experimental mice were co-housed with up to two mice of the same sex per cage with autoclaved standard chow pellets and water provided *ad libitum*. Cages of female mice were treated at 6 weeks of age with an antibiotic cocktail VNA [1 g/L ampicillin, 1 g/L neomycin, 0.5 g/L vancomycin (Sigma, St. Louis, MO)] in drinking water for 7 days.

For fecal and cecal transplants, donor mice were euthanized by CO₂ asphyxiation, and dissections were performed under sterile conditions in a biosafety cabinet using Clidox- and ethanol-sterilized tools. Fecal pellets or cecal contents were collected into sterile 50 mL conical tubes and homogenized in phosphate-buffered saline (PBS). Resulting homogenates were administered to recipient mice by oral gavage using sterile syringes and 20-gauge curved gavage needles on day 8 and day 9, with each mouse receiving approximately 200– 400 µL. For co-housing, Jackson recipient mice were paired with Taconic donor mice on Day 7. Co-housed pairs were maintained in fresh cages with autoclaved bedding, food, and water. Each experimental replicate included control groups: untreated JAX and TAC mice and antibiotic-treated JAX mice with no subsequent transplantation, beyond the antibiotic-treated groups undergoing fecal transplant, cecal transplant, or co-housing. Fecal samples were collected longitudinally, and all mice were monitored daily for behavior, health status, and cage conditions. Special care was taken to prevent cage flooding and ensure adequate chow and water access, particularly during antibiotic treatment. After transplantation or co-housing, all mice received autoclaved drinking water without antibiotics for the remainder of the experiment. All animal procedures were approved by and conducted in compliance with Washington University’s Institutional Animal Care and Use Committee (IACUC) guidelines. The experiment was repeated for a total of three independent experiments, with data combined in subsequent analyses.

### 16S rRNA gene Illumina sequencing

Phenol:chloroform-extracted DNA from fecal pellets was used for both 16S rRNA gene qPCR and sequencing. SYBR green qPCR for the 16S rRNA gene was performed with 515F (5’-GTGCCAGCMGCCGCGGTAA-3’) and 805R (5’-GACTACCAGGGTATCTAATCC-3’) primers to detect the V4 hypervariable region. Samples were prepared in three batches with samples from across experiments and treatment groups semi-randomized, in a supervised manner, between the pools.

Primer selection and PCRs were performed as described previously ^14^. Briefly, each sample was amplified in triplicate with Golay-barcoded primers specific for the V4 region (F515/R806), combined, and confirmed by gel electrophoresis. PCR reactions contained 18.8μL RNase/DNase-free water, 2.5μL 10X High Fidelity PCR Buffer (Invitrogen), 0.5μL 10 mM dNTPs, 1μL 50 mM MgSO4, 0.5μL each of the forward and reverse primers (10 μM final concentration), 0.1μL Platinum High Fidelity Taq (Invitrogen) and 1.0μL genomic DNA. Reactions were held at 94°C for 2 min to denature the DNA, with amplification proceeding for 26 cycles at 94°C for 15s, 50°C for 30s, and 68°C for 30s; a final extension of 2 min at 68°C was added to ensure complete amplification. Amplicons were pooled and purified with 0.6x Agencourt Ampure XP beads (Beckman-Coulter) according to the manufacturer’s instructions. The final pooled samples, along with aliquots of the three sequencing primers, were sent to the Center for Genome Sciences (Washington University School of Medicine) for sequencing by the 2x250bp protocol with the Illumina MiSeq platform.

The resulting FASTQ files were processed using the DADA2 pipeline (v1.28.0) implemented in Nephele (version 2024_Jan_16, commit: 71b6a72)^32^. Initial quality assessment was conducted to determine appropriate truncation and filtering parameters. Reads were trimmed to remove low-quality bases and sequencing primers and truncated based on quality scores to ensure high-confidence base calls^45,46^. Specifically, reads were truncated at the position where the quality score dropped below a threshold (default: 4)^47,48^. Reads with expected errors exceeding a set threshold (default: 5) were discarded to further enhance data quality. Forward and reverse reads were merged, requiring a minimum overlap of 12bp and allowing for zero mismatches in the overlapping region to ensure accurate reconstruction of the target amplicon^49,50^. Chimeric sequences were identified and removed using the consensus method to prevent false-positive operational taxonomic units (OTUs).

Taxonomic classification was performed using the Ribosomal Database Project (RDP) classifier, with a minimum bootstrap confidence threshold of 40 for taxonomic assignments and the SILVA v138.1 database^51^. Additionally, a phylogenetic tree was constructed using MAFFT for multiple sequence alignment and FastTree2 for tree building, facilitating downstream phylogenetic analyses^52,53^. The output included an amplicon sequence variant (ASV) table, taxonomic assignments, and phylogenetic relationships, providing a comprehensive dataset for subsequent diversity and composition analyses. All analyses were conducted using default parameters unless otherwise specified, adhering to best practices for 16S rRNA gene sequence processing to ensure accurate and reproducible results. One sample (“TACNoT.MTP2.D11.74”) was excluded due to lack of stool at the time of DNA extraction. Twelve ASVs assigned to Eukaryota, Mitochondria, or Chloroplast were removed, and one sample that no longer contained bacterial reads were excluded from downstream analysis.

### Bacterial ecological and distance analyses

Bacterial community composition and ecological metrics were assessed using both alpha and beta diversity analyses. Alpha diversity metrics, including observed richness and Shannon diversity, were calculated to evaluate within-sample diversity. Beta diversity was assessed using unweighted and weighted UniFrac distances to capture phylogenetic differences between samples. These analyses were implemented using the ‘phyloseq’ and ‘vegan’ R packages, which offer comprehensive tools for ecological and diversity analyses. Principal coordinate analysis (PCoA) was performed to visualize patterns in beta diversity across treatment groups and time points^54,55^. Pairwise comparisons between groups were conducted using PERMANOVA with Bonferroni corrections to identify statistically significant differences (alpha = 0.05). These analyses provided insights into the ecological shifts in bacterial communities associated with different transplantation methods and recovery stages. All figures were generated in R using ggplot2^56^.

### Identifying bacterial taxa associated with each vendor

Bacterial taxa specific to each vendor were identified using ASVs derived from 16S rRNA gene sequencing. Taxa were classified as Jackson-associated or Taconic-associated based on their differential prevalence and abundance in fecal samples collected from untreated mice during the first week of the experiment (days 0-7). Samples were limited to those without antibiotic treatment to accurately capture baseline community differences. Differential abundance analyses were conducted using the DESeq2 pipeline, with ASVs assigned taxonomy using the SILVA database. Significant differences were determined using an adjusted p-value threshold of 0.01 and a combined log2(fold change) cutoff of ±4. Prevalence was calculated as the proportion of samples in which a given ASV was detected within each vendor group. To incorporate abundance and prevalence, a combined fold-change metric was calculated by summing the log2(fold change) from differential abundance analysis and the log2(prevalence ratio) between the two groups. Statistical and prevalence-based analyses were compared to confirm associations, and taxa that did not meet these criteria were categorized as non-supplier-specific.

### Predicting functional capacity of bacterial communities

Functional capacities of bacterial communities were inferred from 16S rRNA gene sequencing data using the PICRUSt2 (v. 2.5.3) pipeline. ASVs were mapped to predicted genes and metabolic pathways using the MetaCyc database. Differentially abundant pathways were identified using DESeq2 with an alpha threshold of 0.05. Comparisons were made between antibiotic-treated recipient mice (co-housed, fecal transplantation, or cecal transplantation groups) and control communities from untreated Jackson and Taconic mice. Functional convergence over time was assessed across treatment groups to evaluate the recovery of microbial metabolic potential.

### Taxonomic changes following transplantation

Changes in bacterial taxonomic composition were assessed longitudinally using 16S rRNA gene sequencing. ASVs were classified into taxonomic families, and relative abundances were tracked over time in recipient mice across different transplantation methods (co-housing, fecal, and cecal). Differentially enriched taxa were identified using ANCOM-BC2 (v. 2.6.0), and significant changes were determined using Kruskal-Wallis tests with Bonferroni corrections for multiple comparisons (alpha = 0.05). Taxonomic dynamics were visualized using stacked bar plots and principal coordinate analyses (PCoA) of unweighted UniFrac distances.

### Statistical comparisons across gastrointestinal sites

To evaluate differences in observed species richness across experimental groups, we fit a linear mixed-effects model using the lmer function from the lme4 package (v. 1.1-35.5), with Sample Type (gastrointestinal site) and Condition (transplantation method) specified as fixed effects and Sequencing Library included as a random effect to account for variation across sequencing batches^57^. ANOVA as implemented in the lmerTest package (3.1-3) was used to test the significance of fixed effects^58^. Post hoc comparisons of estimated marginal means were conducted using the lsmeans function withing the emmeanspackage (v. 1.10.3) with Tukey’s HSD correction for multiple testing^59^. Group-level differences were visualized using compact letter display from the multcompView package (v. 0.1-10), where groups not sharing a letter are significantly different at α = 0.05^60^.

### Virome Illumina sequencing and analysis

Viral communities were characterized using short-read Illumina sequencing of viral-like particles (VLPs) extracted from fecal samples. DNA was purified from VLPs, and libraries were prepared using standard protocols. Sequencing was performed on an Illumina platform, generating paired-end reads. Reads were assembled into contigs using Flye, which is optimized for accurate assembly of viral genomes from metagenomic data. Viral contigs were identified using stringent criteria for viral hallmark genes and annotated taxonomically using Cenote-Taker2, a tool designed for comprehensive identification and annotation of eukaryotic and prokaryotic viruses.

To account for batch effects associated with preparation of viral sequencing pools, raw viral count tables were first converted to a DESeq2^36^ dataset using the phyloseq_to_deseq2^55^ function with a design formula including VP, Source, Day, Antibiotics, and PostFMT. Counts were log-transformed using variance-stabilizing transformation (VST) with blind = FALSE. Batch effects attributable to viral preparation were then removed using the removeBatchEffect function from the limma package, specifying the model design to preserve variation due to biological covariates^61^. The resulting batch-corrected abundances were rounded to the nearest integer, and any negative values were set to zero before being re-integrated into the phyloseq object for downstream analysis. Samples where fewer than ten unique viral contigs were detected were filtered out of subsequent analyses.

### Viral ecological and distance analyses

Viral community composition and ecological dynamics were assessed using alpha diversity metrics (e.g., richness) and beta diversity measures (e.g., Jaccard distances). These analyses were implemented using the ‘phyloseq’ and ‘vegan’ R packages, which provide robust tools for microbial ecological analyses. Principal coordinate analysis (PCoA) was conducted to visualize temporal changes in viral community structure across treatment groups. Pairwise comparisons between groups were performed using PERMANOVA with Bonferroni corrections for multiple comparisons (alpha = 0.05). Viral communities were further classified as Jackson-associated, Taconic-associated, or non-supplier-specific based on their prevalence and abundance in control communities. Ecological metrics were integrated with bacterial data to explore host-virus interactions during microbiota transplantation. Source discriminatory viral contigs were defined using the combined fold-change metric described for bacteria above.

### Hierarchical clustering of viral contigs

We employed a hierarchical clustering approach to categorize viral contigs into taxonomic units based on amino acid identity (AAI) and average nucleotide identity (ANI). Initial protein-coding genes were predicted from input contigs using Prodigal^62^ (v2.6.3; in metagenomic mode. All-versus-all BLASTP alignments (BLAST+^63^, v2.7.1) were performed to compute AAI values, filtered to retain pairs with ≥20% AAI, ≥10% shared genes, and ≥8 homologous proteins. These filtered edges underwent Markov Clustering (MCL^64^ v22.282) with an inflation parameter of 1.2 to define family-level clusters. Each family cluster was recursively processed using stricter thresholds (≥40% AAI, ≥20% shared genes, ≥16 homologs; MCL inflation=2.0) to resolve genus-level clusters. Finally, genus clusters were subjected to nucleotide-level ANI analysis via BLASTn and checkV’s anicalc.py (checkV^65^ v1.0.1), with vOTUs (viral operational taxonomic units) delineated at ≥95% ANI and ≥85% coverage.

### Viral classification and binning

To further identify viral contigs, we then applied a two-tier filtering approach to these binned contigs. First, any contig shorter than 1,500 bp was excluded. Contigs were then assessed with VirSorter2 (retaining those with a viral score ≥50) and DeepVirFinder (retaining those with a score ≥0.7 and p-value ≤0.05)^66,67^. Contigs that did not meet these primary criteria were retained if they contained more viral genes than host genes and harbored at least one viral hallmark gene as identified by Cenote Taker2^68^. This pipeline yielded 2,262 putative viral contigs. Taxonomic assignments of these viral contigs were determined using vContact3, while iPHoP and PHATyp were employed for host and lifestyle (temperate vs. virulent) predictions, respectively^38,69^. CheckV was used to assess genome quality and completeness^65^. Contigs were binned using CoCoNet, which discards sequences shorter than 2,048 base pairs by deafult^37^.

Bins were classified based on the aggregated predictions of the contigs they contained. Bins were labeled by their inferred source community if they contained any contigs from one of the sources—Taconic-associated or Jackson-associated—and labeled “Neither/Both” otherwise. No bins contained both Taconic-associated and Jackson-associated contigs. Next, lifestyle predictions (virulent vs. temperate) were consolidated at the bin level by checking whether any contigs in a bin were classified as temperate or virulent, and no contigs within the bin were predicted to be of the other lifestyle. No lifestyle classification was assigned to bins with a mixture of predictions at the contig level. Of the classified bins, 165 were predicted to be temperate while 138 were predicted to be virulent. Finally, host family assignments were integrated by examining the taxonomic predictions for each contig within a bin: bins containing contigs from a single family were assigned that family, bins spanning multiple families in a single phylum were labeled with the phylum followed by “_spp,” and bins spanning multiple phyla were labeled “multi.”

### Use of Large Language Models

Large language models, including Google’s Gemini, OpenAI’ ChatGPT, and Apple Intelligence Writing Tools were used in the preparation of this manuscript. These tools were used to edit text and included the use of the following prompts: “Make concise,” “Make concise while preserving figure callouts.” All text output from these models was subsequently edited and confirmed for scientific accuracy by an author.

## Acknowledgments

We acknowledge all members of the Baldridge laboratory for helpful discussions as well as member of June Round’s laboratory.

## Author contributions

J.S.W performed data analysis generated figures and wrote the manuscript. M.E.S., H.M., S.A., M.H., E.A.K., L.A.S. and L.F. performed the experiments. M.E.S., H.M., S.A., and M.T.B. designed the project. L.A.C.C. and A.R contributed analysis of viral data. M.T.B. contributed to the writing of the manuscript. All authors read and edited the manuscript.

## Inclusion and Ethics

We affirm that this work was conducted in accordance with recognized ethical standards in research and reporting. Contributions were made in line with Nature Portfolio authorship guidelines, and all co-authors have approved this version of the manuscript. We are committed to inclusive and equitable collaboration, and we have actively worked to ensure fair representation and acknowledgment of contributions from all team members, regardless of career stage, identity, or background. Where applicable, trainees involved in the study received formal instruction in responsible conduct of research.

## Data and materials availability

The data from this study are tabulated in the main paper and supplementary materials. All reagents are available from M.T.B. under a material transfer agreement with Washington University. Additionally, all sequencing data generated in this study, including 16S rRNA gene amplicon data and virome shotgun metagenomic data, have been deposited in the European Nucleotide Archive (ENA), under accession number PRJEB55403 and publicly accesible upon acceptance. Data files, metadata, and scripts used for analysis are available at Zenodo (**10.5281/zenodo.15191166)** and will be made publicly accessible upon publication. Mapping between raw files submitted to ENA all relevant metadata are available at Zenodo (**10.5281/zenodo.15191166).**

## Code availability statement

Scripts and code used for analysis are available on Zenodo (**10.5281/zenodo.15191166)** and will be made publicly accessible upon publication.

## References

1. Sommer, F. & Bäckhed, F. The gut microbiota — masters of host development and physiology. Nat. Rev. Microbiol. 11, 227–238 (2013).

2. Round, J. L. & Mazmanian, S. K. The gut microbiota shapes intestinal immune responses during health and disease. Nat. Rev. Immunol. 9, 313–323 (2009).

3. Oliphant, K. & Allen-Vercoe, E. Macronutrient metabolism by the human gut microbiome: major fermentation by-products and their impact on host health. Microbiome 7, 91 (2019).

4. Li, T.-T. et al. Microbiota metabolism of intestinal amino acids impacts host nutrient homeostasis and physiology. Cell Host Microbe 32, 661–675.e10 (2024).

5. Donald, K. & Finlay, B. B. Early-life interactions between the microbiota and immune system: impact on immune system development and atopic disease. Nat. Rev. Immunol. 23, 735–748 (2023).

6. Lambring, C. B. et al. Impact of the Microbiome on the Immune System. Crit. Rev. Immunol. 39, 313–328 (2019).

7. Graham, D. B. & Xavier, R. J. Conditioning of the immune system by the microbiome. Trends Immunol. 44, 499–511 (2023).

8. Villarino, N. F. et al. Composition of the gut microbiota modulates the severity of malaria. Proc. Natl. Acad. Sci. U. S. A. 113, 2235–2240 (2016).

9. Baldridge, M. T. et al. Commensal microbes and interferon-λ determine persistence of enteric murine norovirus infection. Science 347, 266–269 (2015).

10. Winkler, E. S. et al. The Intestinal Microbiome Restricts Alphavirus Infection and Dissemination through a Bile Acid-Type I IFN Signaling Axis. Cell 182, 901–918.e18 (2020).

11. Turnbaugh, P. J. et al. An obesity-associated gut microbiome with increased capacity for energy harvest. Nature 444, 1027–1031 (2006).

12. Ridaura, V. K. et al. Gut microbiota from twins discordant for obesity modulate metabolism in mice. Science 341, 1241214 (2013).

13. Gehrig, J. L. et al. Effects of microbiota-directed foods in gnotobiotic animals and undernourished children. Science 365, eaau4732 (2019).

14. Di Luccia, B. et al. Combined Prebiotic and Microbial Intervention Improves Oral Cholera Vaccination Responses in a Mouse Model of Childhood Undernutrition. Cell Host Microbe 27, 899–908.e5 (2020).

15. Britton, G. J. et al. Defined microbiota transplant restores Th17/RORγt+ regulatory T cell balance in mice colonized with inflammatory bowel disease microbiotas. Proc. Natl. Acad. Sci. U. S. A. 117, 21536–21545 (2020).

16. Britton, G. J. et al. Microbiotas from Humans with Inflammatory Bowel Disease Alter the Balance of Gut Th17 and RORγt+ Regulatory T Cells and Exacerbate Colitis in Mice. Immunity 50, 212–224.e4 (2019).

17. Kennedy, E. A., King, K. Y. & Baldridge, M. T. Mouse Microbiota Models: Comparing Germ-Free Mice and Antibiotics Treatment as Tools for Modifying Gut Bacteria. Front. Physiol. 9, (2018).

18. Witjes, V. M., Boleij, A. & Halffman, W. Reducing versus Embracing Variation as Strategies for Reproducibility: The Microbiome of Laboratory Mice. Animals 10, 2415 (2020).

19. Schloss, P. D. Identifying and Overcoming Threats to Reproducibility, Replicability, Robustness, and Generalizability in Microbiome Research. mBio 9, 10.1128/mbio.00525-18 (2018).

20. Huttenhower, C., Finn, R. D. & McHardy, A. C. Challenges and opportunities in sharing microbiome data and analyses. Nat. Microbiol. 8, 1960–1970 (2023).

21. Moossavi, S., Fehr, K., Khafipour, E. & Azad, M. B. Repeatability and reproducibility assessment in a large-scale population-based microbiota study: case study on human milk microbiota. Microbiome 9, 41 (2021).

22. Zhang, C. et al. Transfer efficiency and impact on disease phenotype of differing methods of gut microbiota transfer. Sci. Rep. 12, 19621 (2022).

23. Ericsson, A. C. & Franklin, C. L. Manipulating the Gut Microbiota: Methods and Challenges. ILAR J. 56, 205–217 (2015).

24. Borin, J. M. et al. Fecal virome transplantation is sufficient to alter fecal microbiota and drive lean and obese body phenotypes in mice. Gut Microbes 15, 2236750 (2023).

25. Rasmussen, T. S. et al. Faecal virome transplantation decreases symptoms of type 2 diabetes and obesity in a murine model. Gut 69, 2122–2130 (2020).

26. Federici, S. et al. Targeted suppression of human IBD-associated gut microbiota commensals by phage consortia for treatment of intestinal inflammation. Cell 185, 2879–2898.e24 (2022).

27. Gogokhia, L. et al. Expansion of Bacteriophages Is Linked to Aggravated Intestinal Inflammation and Colitis. Cell Host Microbe 25, 285–299.e8 (2019).

28. Hsu, B. B. et al. Dynamic Modulation of the Gut Microbiota and Metabolome by Bacteriophages in a Mouse Model. Cell Host Microbe 25, 803–814.e5 (2019).

29. Pfeiffer, J. K. & Virgin, H. W. Transkingdom control of viral infection and immunity in the mammalian intestine. Science 351, aad5872 (2016).

30. Brunse, A. et al. Fecal filtrate transplantation protects against necrotizing enterocolitis. ISME J. 16, 686–694 (2022).

31. Caruso, R., Ono, M., Bunker, M. E., Núñez, G. & Inohara, N. Dynamic and Asymmetric Changes of the Microbial Communities after Cohousing in Laboratory Mice. Cell Rep. 27, 3401–3412.e3 (2019).

32. Weber, N. et al. Nephele: a cloud platform for simplified, standardized and reproducible microbiome data analysis. Bioinforma. Oxf. Engl. 34, 1411–1413 (2018).

33. Callahan, B. J. et al. DADA2: High-resolution sample inference from Illumina amplicon data. Nat. Methods 13, 581–583 (2016).

34. Douglas, G. M. et al. PICRUSt2 for prediction of metagenome functions. Nat. Biotechnol. 38, 685–688 (2020).

35. Caspi, R. et al. The MetaCyc database of metabolic pathways and enzymes and the BioCyc collection of pathway/genome databases. Nucleic Acids Res. 44, D471–480 (2016).

36. Love, M. I., Huber, W. & Anders, S. Moderated estimation of fold change and dispersion for RNA-seq data with DESeq2. Genome Biol. 15, 550 (2014).

37. Arisdakessian, C. G., Nigro, O. D., Steward, G. F., Poisson, G. & Belcaid, M. CoCoNet: an efficient deep learning tool for viral metagenome binning. Bioinformatics 37, 2803–2810 (2021).

38. Roux, S. et al. iPHoP: An integrated machine learning framework to maximize host prediction for metagenome-derived viruses of archaea and bacteria. PLOS Biol. 21, e3002083 (2023).

39. Robertson, S. J. et al. Comparison of Co-housing and Littermate Methods for Microbiota Standardization in Mouse Models. Cell Rep. 27, 1910–1919.e2 (2019).

40. Lundberg, R. et al. Human microbiota-transplanted C57BL/6 mice and offspring display reduced establishment of key bacteria and reduced immune stimulation compared to mouse microbiota-transplantation. Sci. Rep. 10, 7805 (2020).

41. Hollister, E. B., Gao, C. & Versalovic, J. Compositional and Functional Features of the Gastrointestinal Microbiome and Their Effects on Human Health. Gastroenterology 146, 1449–1458 (2014).

42. Mu, C., Yang, Y., Su, Y., Zoetendal, E. G. & Zhu, W. Differences in Microbiota Membership along the Gastrointestinal Tract of Piglets and Their Differential Alterations Following an Early-Life Antibiotic Intervention. Front. Microbiol. 8, 797 (2017).

43. Shkoporov, A. N. et al. Viral biogeography of the mammalian gut and parenchymal organs. Nat. Microbiol. 7, 1301–1311 (2022).

44. Anders, J. L. et al. Comparing the gut microbiome along the gastrointestinal tract of three sympatric species of wild rodents. Sci. Rep. 11, 19929 (2021).

45. Andrews, S. Babraham Bioinformatics - FastQC A Quality Control tool for High Throughput Sequence Data. https://www.bioinformatics.babraham.ac.uk/projects/fastqc/.

46. Ewels, P., Magnusson, M., Lundin, S. & Käller, M. MultiQC: summarize analysis results for multiple tools and samples in a single report. Bioinforma. Oxf. Engl. 32, 3047–3048 (2016).

47. Martin, M. Cutadapt removes adapter sequences from high-throughput sequencing reads. EMBnet.journal 17, 10–12 (2011).

48. Bolger, A. M., Lohse, M. & Usadel, B. Trimmomatic: a flexible trimmer for Illumina sequence data. Bioinformatics 30, 2114–2120 (2014).

49. Magoč, T. & Salzberg, S. L. FLASH: fast length adjustment of short reads to improve genome assemblies. Bioinformatics 27, 2957–2963 (2011).

50. Streett, D. dstreett/FLASH2. (2025).

51. Quast, C. et al. The SILVA ribosomal RNA gene database project: improved data processing and web-based tools. Nucleic Acids Res. 41, D590–D596 (2013).

52. Katoh, K. & Standley, D. M. MAFFT Multiple Sequence Alignment Software Version 7: Improvements in Performance and Usability. Mol. Biol. Evol. 30, 772–780 (2013).

53. Price, M. N., Dehal, P. S. & Arkin, A. P. FastTree 2 – Approximately Maximum-Likelihood Trees for Large Alignments. PLoS ONE 5, e9490 (2010).

54. Oksanen, J., et al. vegan: Community Ecology Package. 2.6-8 10.32614/CRAN.package.vegan (2001).

55. McMurdie, P. J. & Holmes, S. phyloseq: An R Package for Reproducible Interactive Analysis and Graphics of Microbiome Census Data. PLOS ONE 8, e61217 (2013).

56. Wickham, H., et al. ggplot2: Create Elegant Data Visualisations Using the Grammar of Graphics. (2024).

57. Bates, D., Mächler, M., Bolker, B. & Walker, S. Fitting Linear Mixed-Effects Models Using lme4. J. Stat. Softw. 67, 1–48 (2015).

58. Kuznetsova, A., Brockhoff, P. B. & Christensen, R. H. B. lmerTest Package: Tests in Linear Mixed Effects Models. J. Stat. Softw. 82, 1–26 (2017).

59. Lenth, R. V., et al. emmeans: Estimated Marginal Means, aka Least-Squares Means. (2025).

60. Graves, S., Piepho, H.-P. & Selzer, L. multcompView: Visualizations of Paired Comparisons. (2024).

61. Ritchie, M. E. et al. limma powers differential expression analyses for RNA-sequencing and microarray studies. Nucleic Acids Res. 43, e47 (2015).

62. Hyatt, D. et al. Prodigal: prokaryotic gene recognition and translation initiation site identification. BMC Bioinformatics 11, 119 (2010).

63. Camacho, C. et al. BLAST+: architecture and applications. BMC Bioinformatics 10, 421 (2009).

64. Van Dongen, S. Graph Clustering Via a Discrete Uncoupling Process. SIAM J. Matrix Anal. Appl. 30, 121–141 (2008).

65. Nayfach, S. et al. CheckV assesses the quality and completeness of metagenome-assembled viral genomes. Nat. Biotechnol. 39, 578–585 (2021).

66. Guo, J. et al. VirSorter2: a multi-classifier, expert-guided approach to detect diverse DNA and RNA viruses. Microbiome 9, 37 (2021).

67. Ren, J. et al. Identifying viruses from metagenomic data using deep learning. Quant. Biol. 8, 64–77 (2020).

68. Tisza, M. J., Belford, A. K., Domínguez-Huerta, G., Bolduc, B. & Buck, C. B. Cenote-Taker 2 democratizes virus discovery and sequence annotation. Virus Evol. 7, veaa100 (2021).

69. Shang, J., Tang, X. & Sun, Y. PhaTYP: predicting the lifestyle for bacteriophages using BERT. Brief. Bioinform. 24, bbac487 (2023).

