## Extended Data Figures for "Dynamics of Bacterial and Viral Transmission in Experimental Microbiota Transplantation"

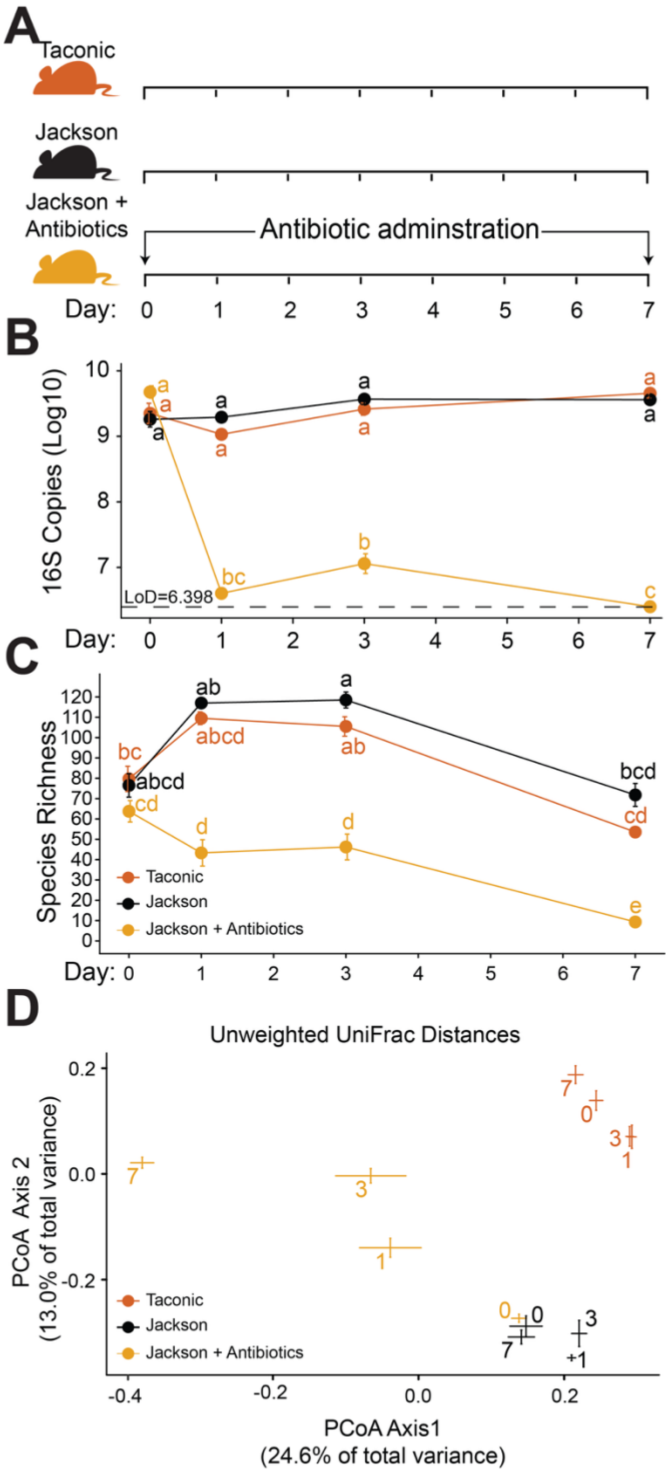


**Extended Data Figure 1. One-week treatment with antibiotic cocktail significantly alters fecal bacterial biomass and community composition.** (**A**) Mice were obtained from Jackson or Taconic labs. Mice from Jackson Labs were provided vancomycin, neomycin, and ampicillin in their drinking water for one week (Jackson + Antibiotics; n=13, 16, and 16 in three independent experiments). Untreated Jackson (n=~4 per experiment each) control mice and Taconic (n=~4 per experiment) mice were maintained on normal drinking water. (**B**) Bacterial load measured by qPCR of 16s rDNA gene throughout the first week of the experiment. The letters next to each point indicate significant groupings, as those with the same letter were not significantly different from one another by two-way ANOVA followed by Tukey's HSD post-hoc test for pairwise comparisons with an alpha of 0.05. (**C**) Bacterial species richness, the number of unique bacteria species observed per mouse, throughout the first week of the experiment via V4-16S rRNA gene sequencing. The letters next to each point indicate significant groupings, as those with the same letter were not significantly different from one another by Kruskal-Wallis followed by Bonferroni's correction for multiple comparisons with an alpha threshold of 0.05. (**D**) Principal Coordinate Analysis of unweighted UniFrac distances calculated from species counts. Error bars (± SE) intersect at centroid of all samples collected for each group of mice on the day post-antibiotics treatment, indicated by the number on the plot.

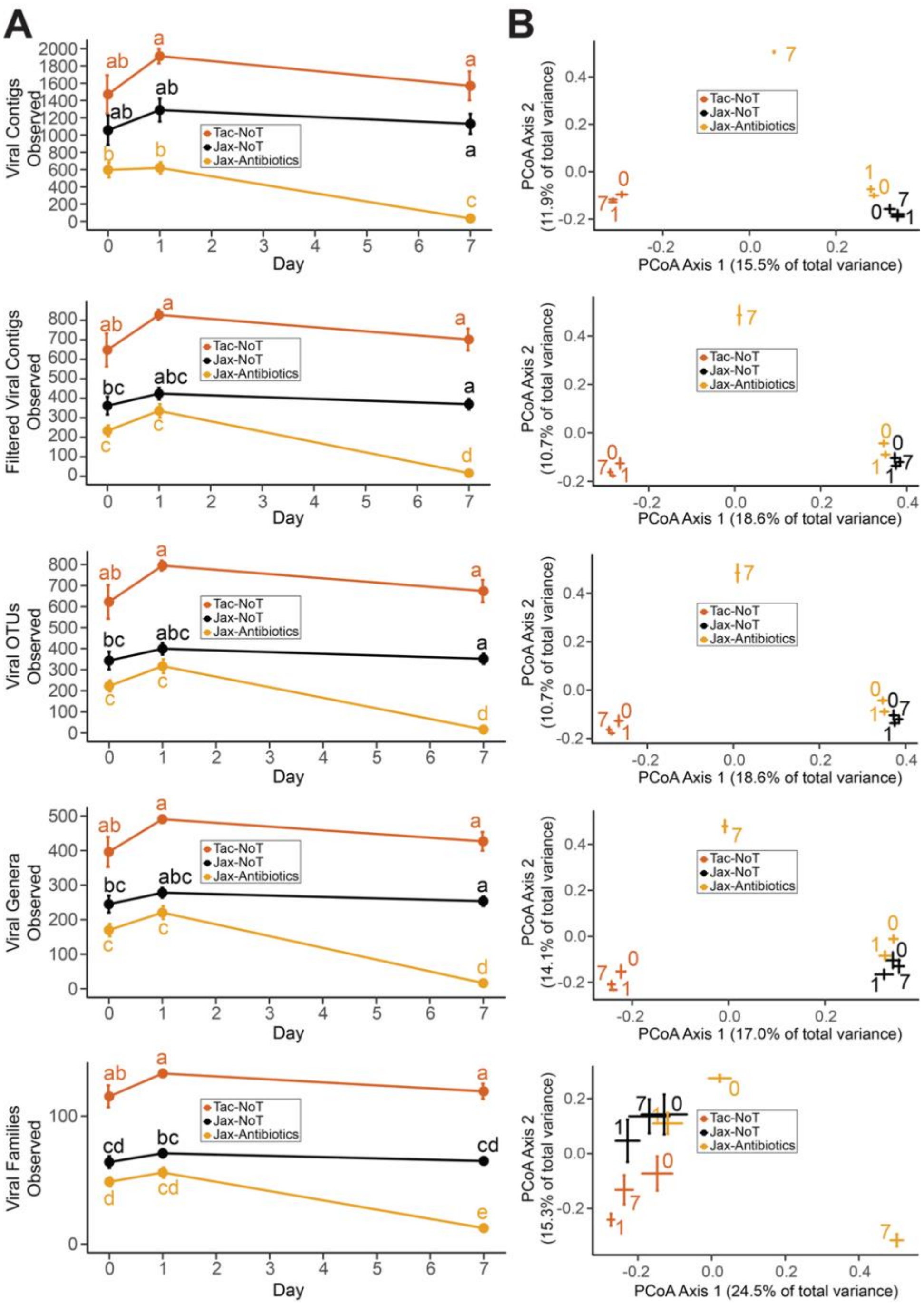

**Extended Data Figure 2. One-week treatment with antibiotic cocktail significantly alters fecal viral richness and community composition.** (**A**) Viral richness and (**B**) Principal Coordinate Analysis of Jaccard distances calculated from contig read counts of unique viral contigs observed per mouse, throughout the first week of the experiment via shotgun metagenomics sequencing of viral-like particles from feces. Panels represent data before (i) and after contigs had been stringently filtered and grouped into clusters representing ii) strains, iii) species/OTUs, iv) genera, and v) families. In A, the dot indicates the mean richness for that group, error bars (± SE), in B, the error bars (± SE) intersect at the centroid of all samples collected for each group of mice on the day indicated by the number on the plot. In A, the letters next to each point indicate significant groupings, those with the same letter were not significantly different from one another by pairwise Kruskal-Wallis followed by Bonferroni’s correction for multiple comparisons with an alpha threshold of 0.05. n = 3-8 samples per treatment group and timepoint.
